# Angular Deviation Diffuser: A Transformer-Based Diffusion Model for Efficient Protein Conformational Ensemble Generation

**DOI:** 10.1101/2025.03.03.640492

**Authors:** Yi Yang, ChuanYe Xiong, Peng Tao

## Abstract

Protein functionality is inherently tied to its structure, with both static and dynamic conformations playing critical roles in defining biological activity. While molecular dynamics (MD) simulations have long been the standard for exploring protein dynamics, they come with high computational costs and limited sampling efficiency. Recent advances in deep learning, such as AlphaFold, have significantly improved static protein structure prediction, yet accurately generating the dynamic ensemble of protein conformations remains a complex challenge. In this study, we present a transformer-based diffusion model that generates diverse conformational ensembles of protein backbones by utilizing angular deviations as data flow. Our model combines a cutting-edge diffusion model with the principles of SE(3) symmetry to enhance both the accuracy and efficiency of conformational sampling. Applied to the Vivid (VVD) Photoreceptor protein system, the generated ensembles closely align with those from MD simulations while covering a broader range of conformational states. This approach offers an improved methodology for capturing protein dynamics, contributing to a more comprehensive understanding of protein structure and function.

## 1. Introduction

The three-dimensional (3D) structure of proteins plays a pivotal role in determining their function within biological systems. This fundamental principle forms the basis of molecular biological systems, as the specific folding and arrangement of amino acid chains into distinct structural motifs(e.g., alpha helices, beta sheets, and loops.) enable proteins to carry out a wide array of biological functions[1]. The dynamics of protein structures, i.e., their conformational flexibility and ability to adopt multiple states, are equally crucial. Therefore, understanding both the static structures and the dynamic conformational landscapes of proteins is essential for elucidating the molecular mechanisms that drive life processes[2].

Traditionally, molecular dynamics (MD) simulations have been the primary tool for exploring the dynamic properties of protein structures. MD simulations provide a time-resolved picture of protein motion by numerically solving Newton’s equations of motion for the atoms within the protein and its surrounding environment. This method offers detailed insights into how proteins explore their conformational space over time[3]. However, one of the major limitations of MD simulations is their requirement for extensive computational resources to capture the full range of possible protein conformations. Achieving adequate sampling often necessitates long simulation times, especially for large or complex systems, resulting in a bottleneck in the efficiency and feasibility of these studies[4]. Despite their accuracy, the low sampling efficiency of MD simulations hinders their ability to capture rare but functionally relevant conformations, thus limiting their utility in generating comprehensive conformational ensembles[5].

To overcome the challenges associated with traditional MD simulations, several works have been developed to enhance or alternative MD approaches. One of the primary strategies involves the use of enhanced sampling techniques, such as *Gaussian accelerated molecular dynamics*[6]*, metadynamics*[7]*, and replica exchange molecular dynamics (REMD)*[8]. These techniques are designed to increase the efficiency of conformational sampling by altering the energy landscape or by simulating multiple trajectories in parallel. Also, LAST technology has been developed by our group in 2023, which introduced a VAE model to enhance the MD sampling process[9]. While these methods have improved the ability to capture diverse protein conformations, they still rely on extensive computational resources and require careful calibration[10].

In parallel with the development of enhanced sampling methods, the rapid advancements in generative artificial intelligence (AI) have opened new avenues for protein conformation generation[11]. Generative AI models, such as *Variational Autoencoders (VAEs)*[12–16]*, Generative Adversarial Networks (GANs)*[17], diffusion model[18–20], and Flow Matching [21] have shown significant promise in producing ensembles of protein conformations that resemble those obtained from MD simulations[22]. These models leverage deep learning techniques to learn the underlying distribution of protein structures from a set of training data and can generate new conformations by sampling from this learned distribution. Despite the significant progress these deep learning-based approaches make, there are some limitations[18]. In particular, the current models struggle to accurately generate conformations for complex protein systems, which are characterized by the presence of small molecules or cofactors that significantly influence the protein’s conformational ensemble, such as vivid Photoreceptor protein system proteins[23,24]. The inclusion of these small molecules complicates the protein’s conformational landscape, making it challenging for existing generative models to produce accurate and diverse ensembles[25,26].

In response to these challenges, and inspired by the groundbreaking work of FoldingDiff[27], we have developed a transformer-based angular deviation diffusion model. Our model introduces several key innovations that enhance the performance of previous approaches. Firstly, it incorporates the SE(3) invariance of protein structures, ensuring that the generated conformations are consistent with the rotational and translational symmetries inherent to molecular systems. Secondly, our model explicitly takes into account the dynamic behavior of proteins as observed in MD simulations by using angular deviation as data flow, allowing it to generate conformations that reflect the dynamics of protein structures. Finally, the diffusion algorithm improves the performance to generate protein backbones.

To validate our model, Firstly, we ran an MD simulation using the VVD protein system and then we trained our model using a slightly short segment of vivid protein simulation conformations. Although limited in length of MD trajectory, this training set provided sufficient data to capture the essential features of the protein’s conformational landscape. Remarkably, our model was able to generate a significantly larger ensemble of conformations in a fraction of the time required for traditional MD simulations. The generated conformations not only closely ensemble those obtained from MD simulations, with a small average RMSD deviation, but also include novel conformations that were not present in the original training set. These results highlight the potential of our transformer-based angular dynamical diffusion model to enhance the efficiency and accuracy of protein conformation generation, particularly for complex systems where previous methods fall short.

## 2. Results

In this study, we first constructed a molecular dynamics (MD) simulation system using the VVD protein system. The simulation was conducted over a span of 1.1 microseconds, and the structural ensemble from the final 1 microsecond of this simulation was utilized as the MD simulation’s conformation set. From this set, we extracted the backbone angle deviations as training data for our model. We then trained a transformer-based diffusion model using a denoising diffusion probabilistic model (DDPM) algorithm[28]. This model was employed to generate a broader and more diverse ensemble of protein backbone structures. Subsequently, we used PyRosetta[29] to add side chains and hydrogen atoms to these backbone structures, followed by further structural optimization, resulting in a complete set of protein conformations (Figure 1).

**Figure 1.**
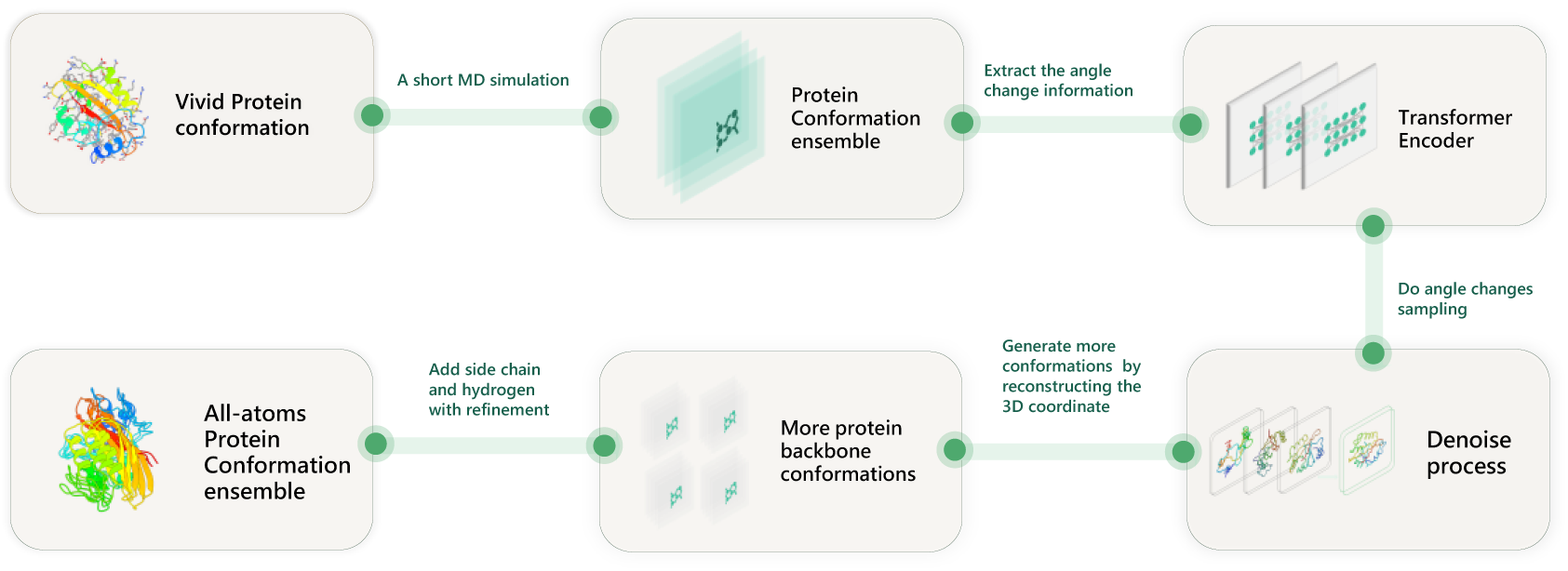
Overview of our work.

### 2.1 Using Angle Deviation is Superior to Using Angles

Inspired by the work of FoldingDiff[27], we extracted six types of angles from the protein backbone structures during the MD simulation. These angles included three types of dihedral torsions (’ϕ’, ‘ψ’, ‘ω’) and three types of bond angles (’θ₁’, ‘θ₂’, ‘θ₃’), which were used as one data point of the training set data for our model (Figure 2). Given the dynamic nature of the structure during MD simulation, and consequently the dynamic nature of the angles, we calculated the angular deviations relative to the structure from the first timestep of the MD conformation set. This angular deviation data was then used as the other training set data. To compare the performance differences generated by using the two different training data types, we conducted ten experiments for each model on two systems (Dark and Light states), generating 500 structures per experiment. We calculated the proportion of correctly folded structures, defined as those with an RMSD (Root Mean Square Deviation) of less than 6 Å compared to the first generated structure, that is the maximum RMSD referred to the first structure generated by MD simulation[25](supplementary figure 1a).

**Figure 2.**
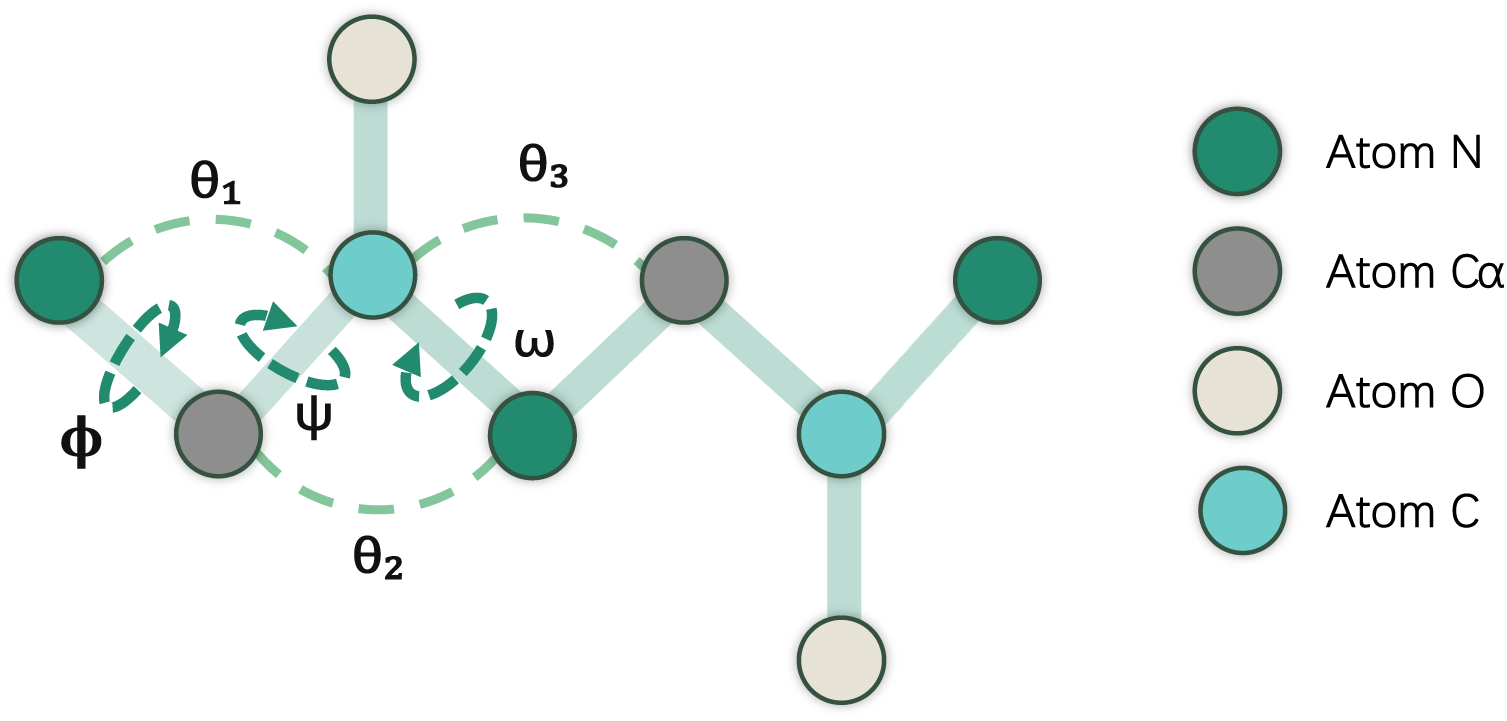
Six angle types of the protein backbone, including bond (light green) and dihedral (dark green) angle.

Our results showed that using angular deviations instead of absolute angles significantly improved the stability of the generated protein structures. For the Dark state system, the average proportion of correctly folded structures was 0.95 when using angular deviations, compared to 0.51 when using absolute angles. Similarly, for the Light state system, the average proportion of correctly folded structures was 0.91 with angular deviations and 0.43 with absolute angles (figure 3a). Moreover, structures generated using angular deviations folded more rapidly and easily during the denoising process, and the angle distributions learned by the model were closer to those observed in the MD simulation. For the Dark state system, the area under the RMSD curve relative to the final folded structure during denoising was smaller when using angular deviations than when using absolute angles. Additionally, the Jensen-Shannon (JS) divergence between the generated angle distributions and the MD simulation angle distributions was lower when using angular deviations (0.13) compared to absolute angles (0.25) (Figure 3b).

**Figure 3.**
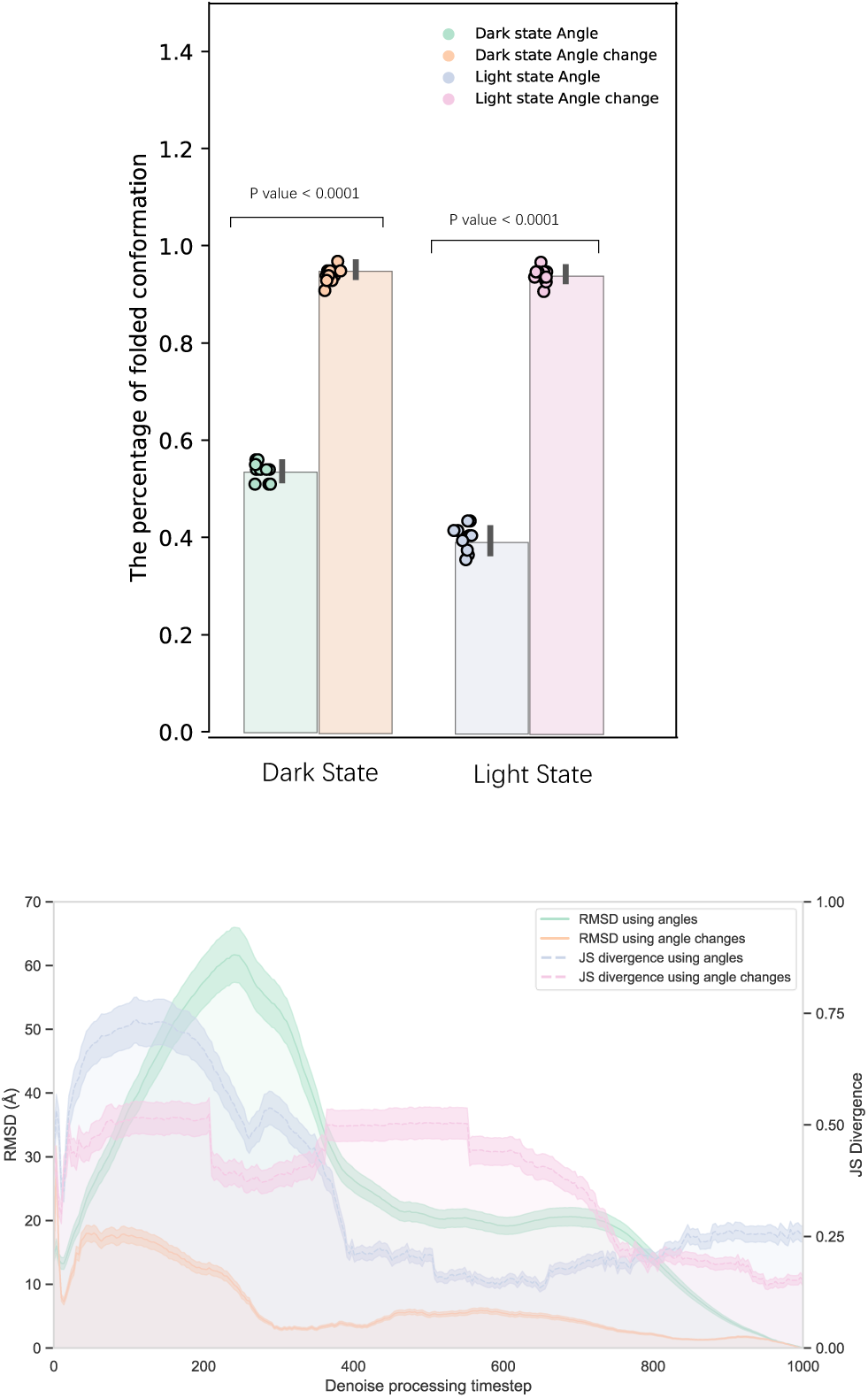
Using angle deviation versus using absolute angle. (a) Comparison of the proportion of correctly folded structures generated using absolute angles versus angular deviation (b) Comparison of RMSD area under the curve and JS divergence between generated angle distributions and MD simulation angle distributions.

### 2.2 Evaluation of the Generated Ensembles

To evaluate the structural ensembles generated by the model, we generated 6000 structures for each of the two systems (Dark state and Light state). First, we compared the distributions of the six backbone angles between the generated structures and those from the MD simulation (training set). The distributions were highly similar, for the dark state, the six-angle average JS divergence was 0.16, and for the light state, the six-angle average JS divergence was 0.15, indicating that the generated ensembles successfully captured the angle distributions of the original MD simulation (Figure 4) for both dark state and light state.

**Figure 4.**
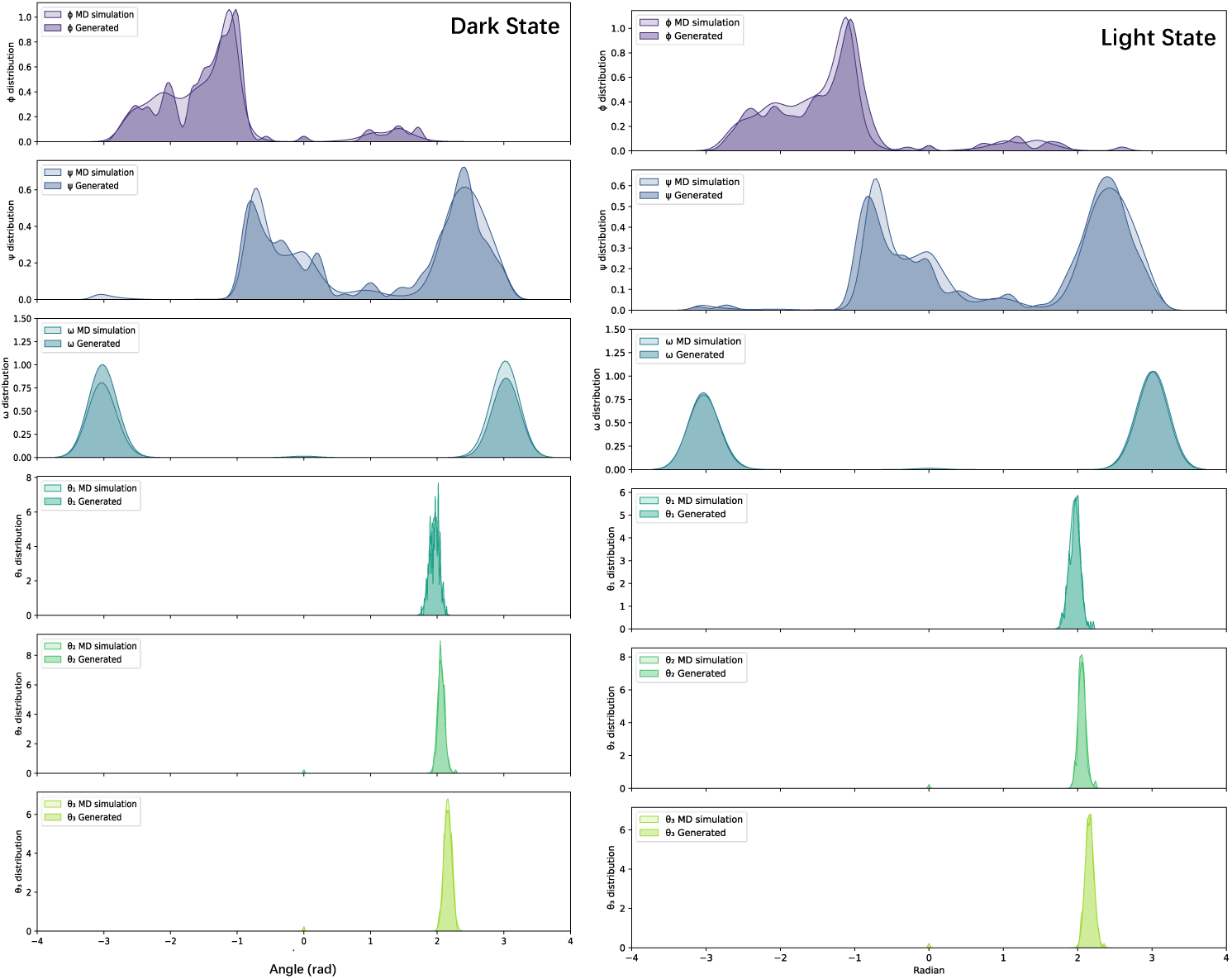
The distributions of the six types of angles between the generated structures and the MD simulation. (a) Dark state system (b) Light state system.

To further analyze the topology of our generated structures, we plotted the Ramachandran diagrams of the generated ensembles, which revealed distinct secondary structure features typical of protein structures, including alpha helix and beta sheet (Figure 5). Based on our Ramachandran diagram results, we found that our generated structures possess reasonable second structures (e.g., Right-handed/Left-handed Helix, Sheet). Moreover, we compared the RMSD distributions of the generated structures with those of the training set. The results showed that our model was able to generate a wider range of structures, with a broader RMSD distribution, indicating the model’s capability to explore a more extensive conformational space (Figure 6).

**Figure 5.**
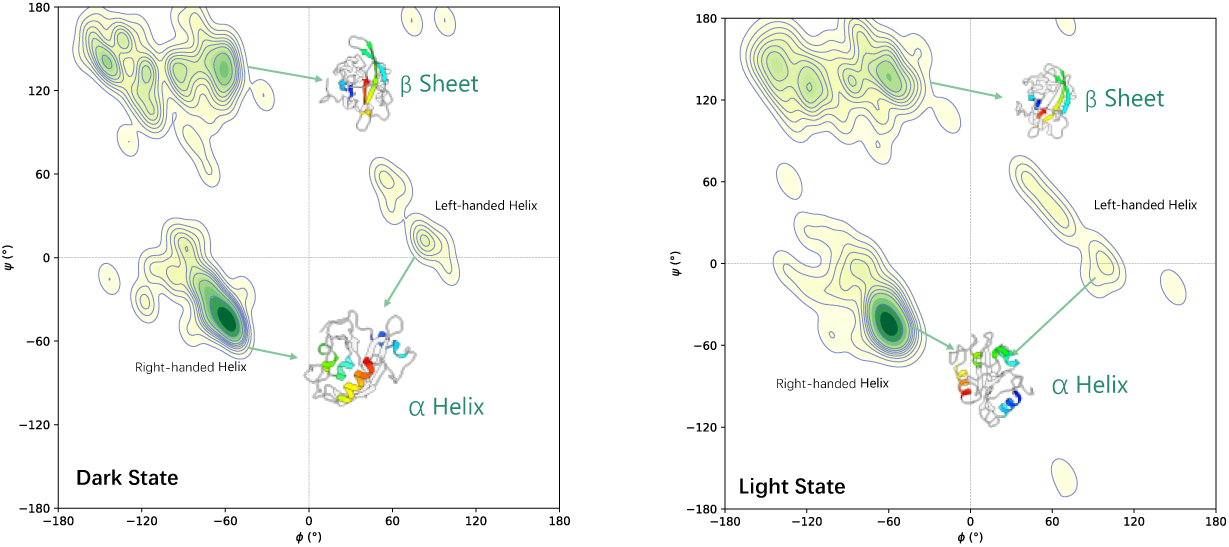
Ramachandran plot of the generated ensembles. (a) dark state system (b) light state system

**Figure 6.**
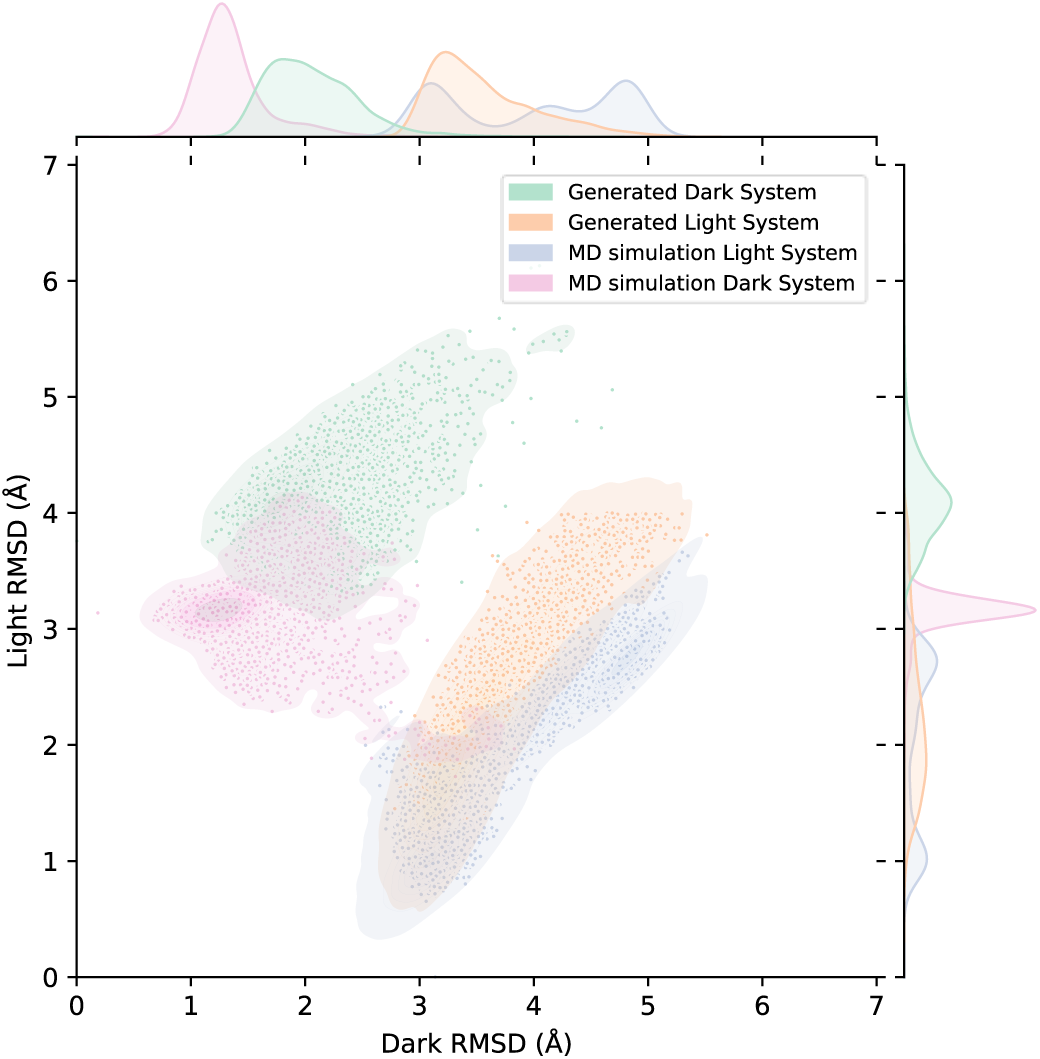
RMSD distributions of the generated structures with the training set. The x-axis represents RMSD for the dark state, while the y-axis represents RMSD for the light state.

### 2.3 Comparison with AlphaFold Flow

To evaluate the performance of our model in comparison to other methods, we generated 6000 structural conformations using our model trained separately on both the Dark and Light state systems of the VVD protein. These structures were subsequently optimized to enhance accuracy. For comparative analysis, we generated an additional set of 6000 structures using AlphaFold Flow by the amino acid sequence of the Dark and Light state system[21]. AlphaFold Flow is a variant of the AlphaFold model tailored for generating molecular dynamics (MD) conformational ensembles. This set was constructed using the same amino acid sequences for the VVD protein in both the Dark and Light states.

We calculated the pairwise RMSD between these four conformational ensembles and the original MD simulation ensemble (training set) of 6000 structures for two systems: the Dark and Light states. In the Dark state, the structures generated by our model with refinement yield a mean RMSD of approximately 3.12 Å, showing better accuracy than the AlphaFold Flow (mean RMSD ∼15.79 Å) and very close to the new MD simulation (mean RMSD ∼1.27 Å)(Figure 7a). In the Light state, the RMSD values show a similar trend, with the refined model having a mean RMSD of 2.54 Å, also very close to the new MD simulation (mean RMSD ∼1.60 Å)(Figure 7b).

**Figure 7:**
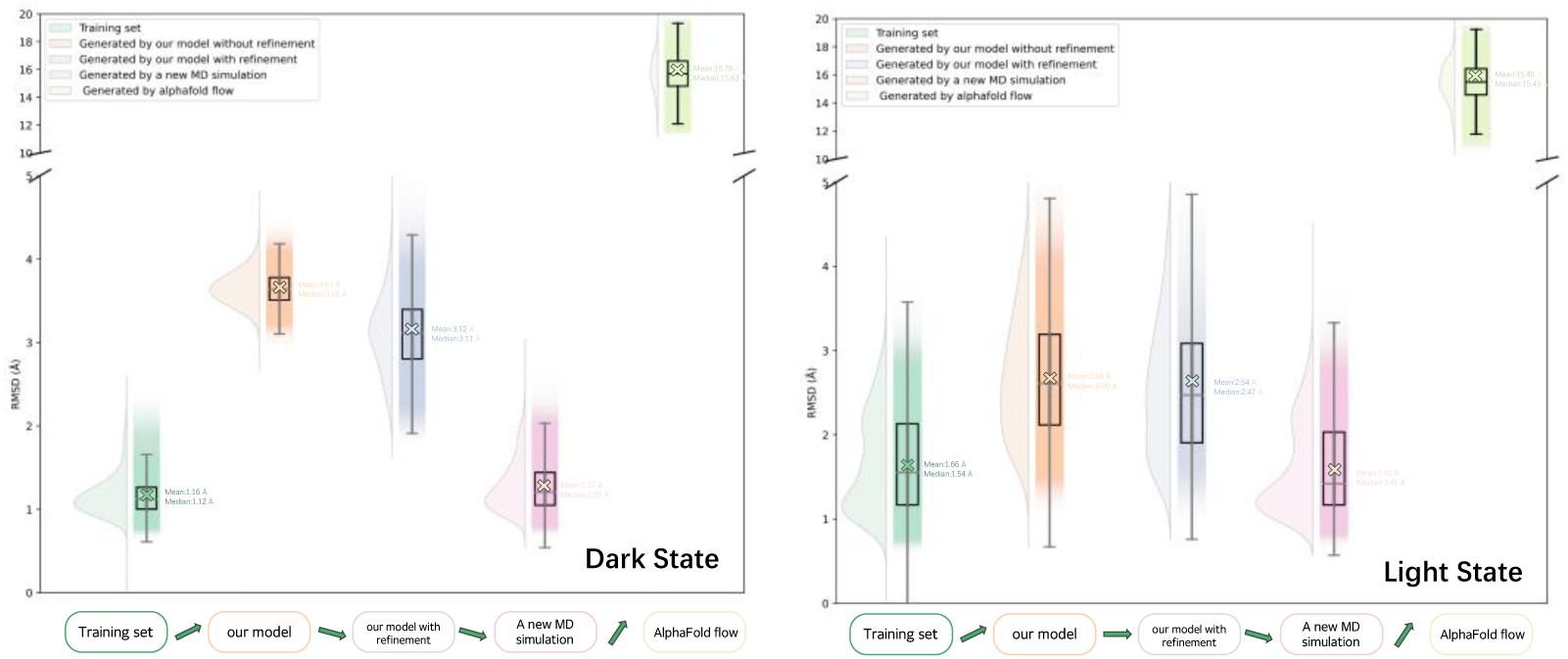
Pairwise RMSD distributions comparing five distinct conformational ensembles. (i) the training set, (ii) structures generated by our model without refinement, (iii) structures generated by our model with refinement, (iv) a new MD simulation(v) structures generated using AlphaFold Flow—against the original MD simulation ensemble(the training set). (a) Pairwise RMSD distributions for the Dark state system; (b) Pairwise RMSD distributions for the Light state system.

These comparisons demonstrate that our model, particularly with refinement, produces structures closer to the original MD ensemble than those generated by AlphaFold Flow, suggesting its robustness in accurately capturing the conformational landscape of the VVD protein in both states.

## 3 Materials and Methods

### 3.1 Molecular Dynamics (MD) Simulation of Vivid (VVD) Photoreceptor protein

Vivid (VVD) is a photoreceptor protein found in the filamentous fungus Neurospora crassa, which plays a crucial role in modulating the organism’s circadian rhythms. VVD contains a light, oxygen, or voltage (LOV) domain, which is a small, highly conserved element responsible for coupling blue-light activation to subsequent protein conformational changes[24,25]. LOV domains are widely present in bacteria, archaea, fungi, and plants, facilitating their blue-light responses. A common feature of all LOV domains is the presence of a flavin cofactor, such as flavin adenine dinucleotide (FAD), flavin mononucleotide (FMN), or riboflavin, which is critical for their function[30].

In this study, we employed molecular dynamics (MD) simulations to investigate the conformational changes in VVD between its dark-adapted and light-activated states. The initial structures for the MD simulations were obtained from the crystal structures of VVD in its dark state (PDB ID: 2PD7) and light state (PDB ID: 3RH8), both of which include the FAD cofactor. To ensure consistency in the sequence lengths between the dark and light states, we removed the 36th residue from the dark state structure. Additionally, given the biological similarity between FMN and FAD, we modified the FAD molecule in the crystal structures by removing the AMP moiety to generate an FMN molecule. The force field parameters for FMN were adopted from previous studies, allowing us to accurately model its behavior within the simulation system[31]. A total of two simulation systems were constructed based on the modified crystal structures (figure 8). Hydrogen atoms were added to each structure, and the systems were solvated using the explicit water model TIP3P[32]. The systems were then neutralized by adding sodium cations and chloride anions. For more details about MD simulation, please see the supplement material.

**Figure 8.**
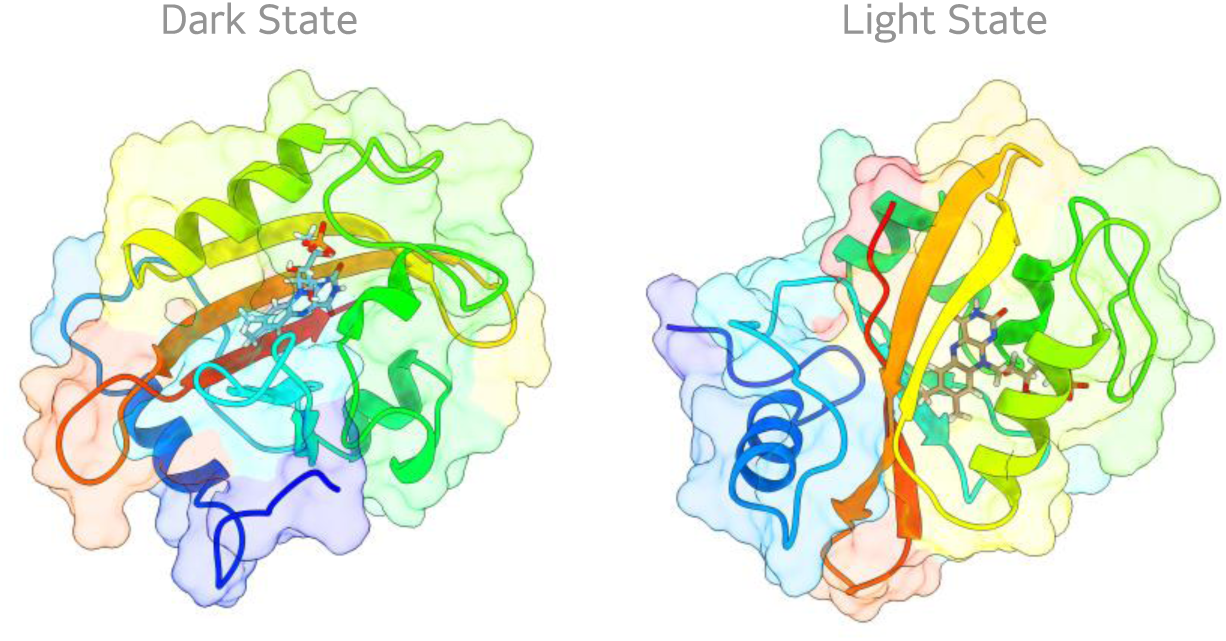
Two MD simulation systems, dark(PDB ID 2pd7) and light(PDB ID 3rh8) state systems.

### 3.2 Diffusion model

The diffusion model framework comprises two main processes: the forward diffusion process and the backward generation process. In the forward process, noise is incrementally added to the protein conformational data by sampling from a wrapped normal distribution[27]. Mathematically, the forward diffusion process is represented by Equation (1):

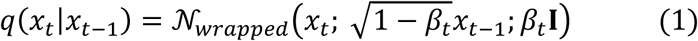

Where I is the identity matrix, *β*_*t*_ is the hyperparameter regarding the noise scheduling. The backward process aims to denoise the input from pure noise to the generated data. The transformer model was utilized to predict the distribution of *p*_*θ*_(*x*_*t*_|*x*_*t*−1_) shown as the equation (2):

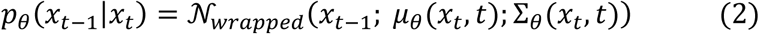

where *μ*_*θ*_(*x*_*t*_, *t*) and Σ_*θ*_(*x*_*t*_, *t*) are the predicted mean value and covariance matrix via the transformer model based on the data at time step t, respectively.

The algorithm of the training process (table 1) is utilized from the previous work[28] where we trained the transformer model by conducting the gradient decent based on loss function *L*_*w*_, and as for the sampling process we modified the noise sampling step shown as equation (3):

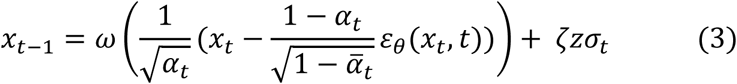

**Table 1.**
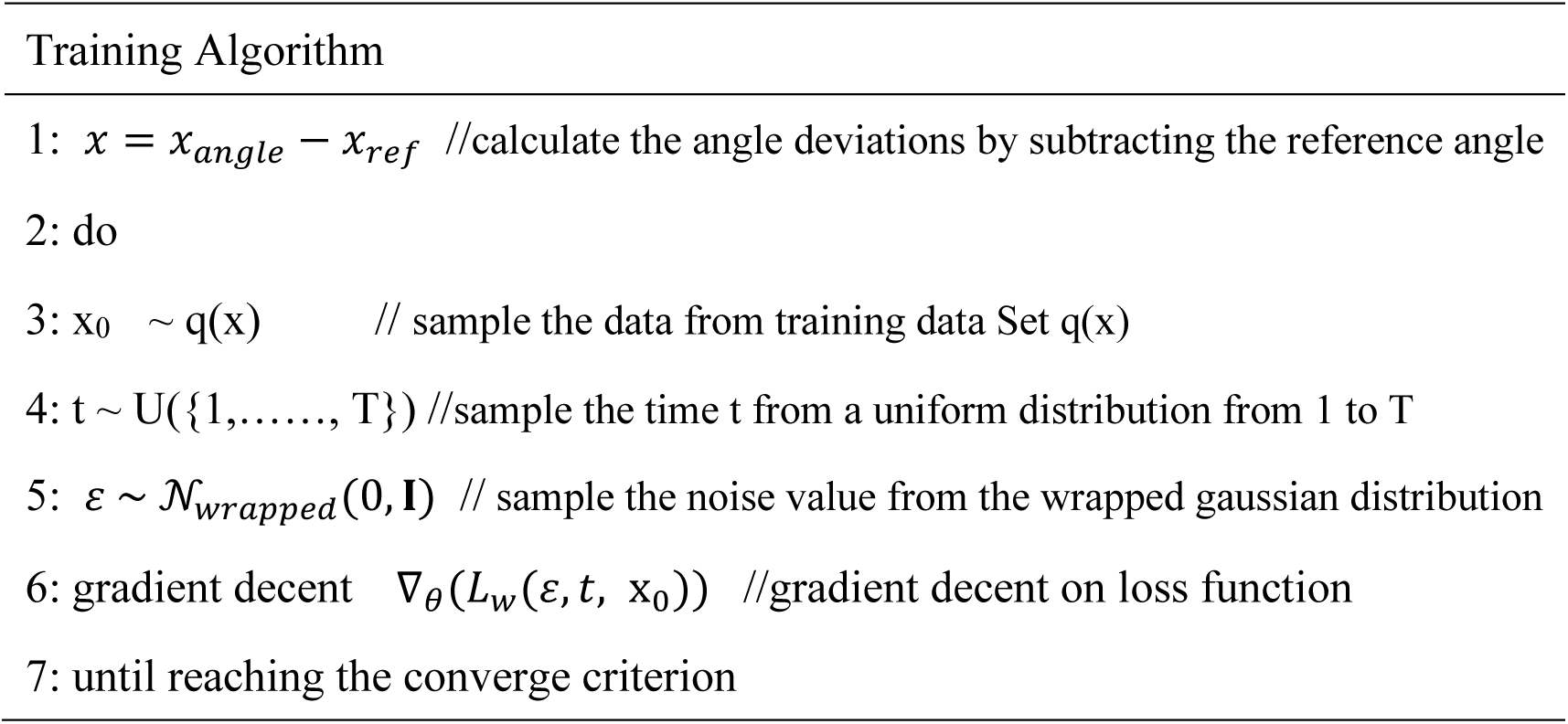
training process.

| Training Algorithm |
| --- |
| 1: $x = x_{angle} - x_{ref}$ //calculate the angle deviations by subtracting the reference angle |
| 2: do |
| 3: $x_0 \sim q(x)$ // sample the data from training data Set $q(x)$ |
| 4: $t \sim U(\{1, \dots, T\})$ //sample the time $t$ from a uniform distribution from 1 to $T$ |
| 5: $\varepsilon \sim \mathcal{N}_{wrapped}(0, \mathbf{I})$ // sample the noise value from the wrapped gaussian distribution |
| 6: gradient decent $\nabla_{\theta}(L_w(\varepsilon, t, x_0))$ //gradient decent on loss function |
| 7: until reaching the converge criterion |

Where the *ω*(*X*) is the wrapping function to achieve the periodic nature of angular data:

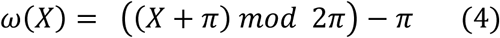

Also, the *ζ* refers to the empirical scaling coefficient to boost the data generation process[16], the empirical value is optimized to be 0.015 in our system. ε_*θ*_(*x*_*t*_, *t*) refers to the predicted noise value by the transformer model at time step t(table 2).

**Table 2.**
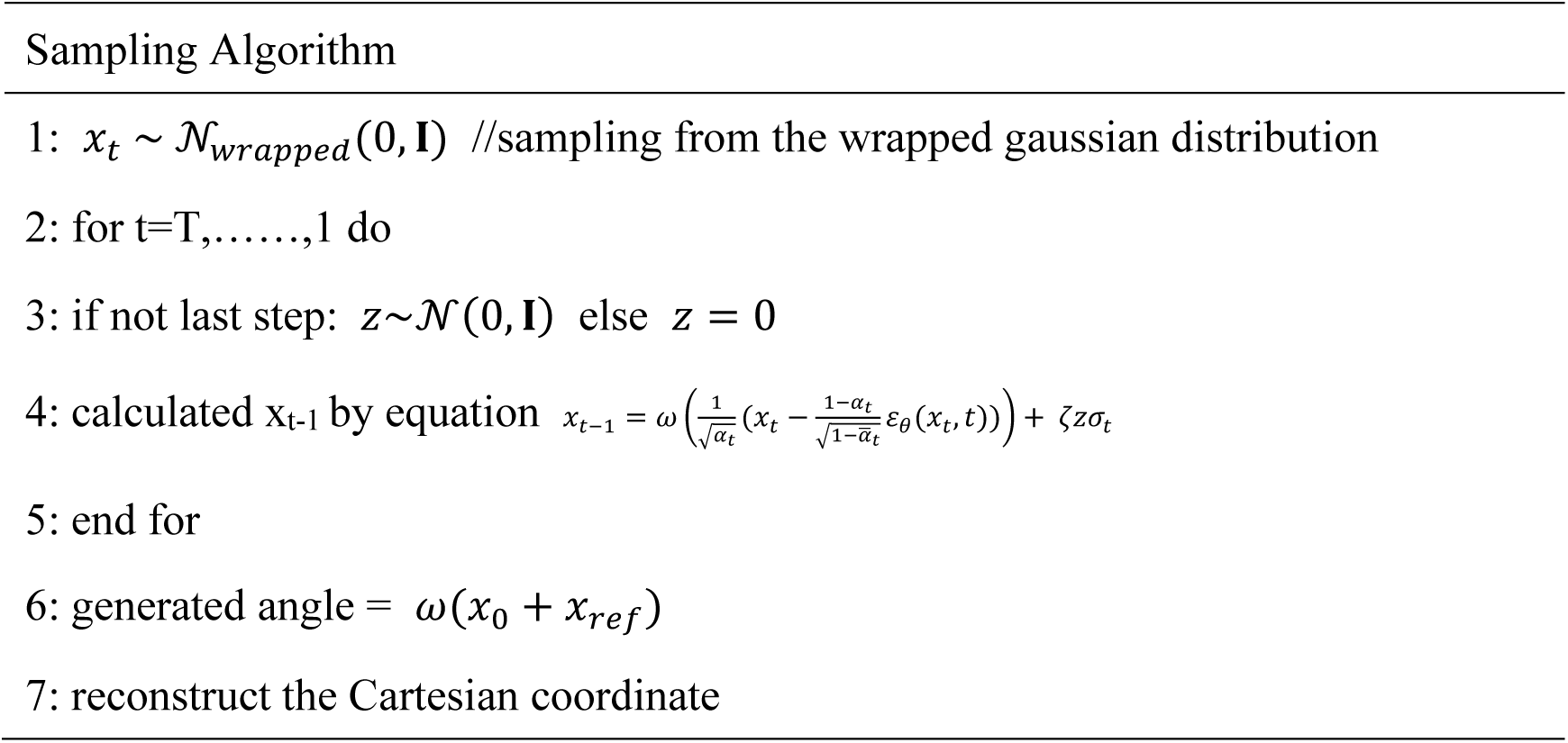
Sampling process.

### 3.3 Transformer Model

To denoise the data, the transformer encoder architecture was adopted[33]. Our structure data were projected to the 147 × 6 *dimension* where 147 is our residue number and 6 is for angle changes, where 3 dihedral angle changes and 3 bond angle changes for each residue. The 6-dimension angle input is projected to the transformer’s 384 embedding dimensions. For more details about our model, please see the supplement material.

### 3.4 Training Set and Loss Function

From molecular dynamics (MD) simulations, we collected a total of 6,001 distinct conformations of the VVD protein of each system. To simplify the analysis, we excluded the small molecule FAD (flavin adenine dinucleotide) from each conformation and extracted the backbone structure for further investigation. Specifically, we computed six backbone angles for each conformation and calculated the changes in these angles relative to a reference structure.

For model training, we utilized 90% of the dataset as the training set, reserving the remaining 10% for validation. We applied a “warm-up” learning rate strategy to ensure stable model convergence[34]. To mitigate the risk of overfitting, we saved model parameters every 200 steps after the model began converging. Finally, we selected the set of model parameters that demonstrated the best performance on the training set as the final parameters for the diffusion model. A wrapped smooth L1 loss was adapted according to the previous literature[27,35], the expression for *L*_*w*_ shown below:

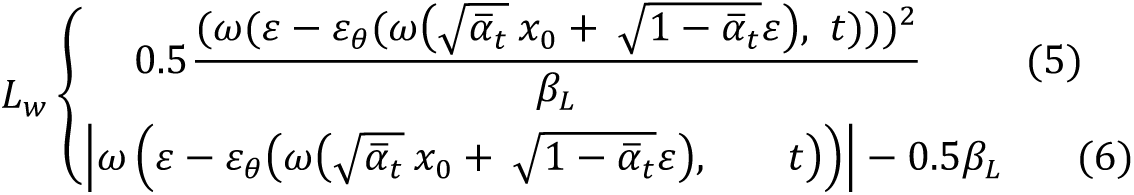

If the 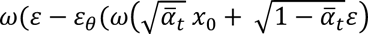 term is smaller than *β*_*L*_, the loss is calculated by equation (5), otherwise loss is calculated by (6). Where *β*_*L*_ is the transition boundary and *β*_*L*_ = 0.1π.

### 3.5 Root Mean Square Deviation (RMSD)

RMSD was calculated by using the equation of (7) to measure the quality of the generated protein structure by comparing the topological differences between the generated protein structures and the output structures from MD calculation.

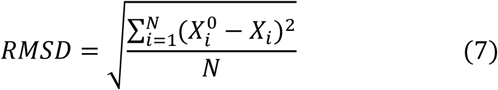

Where 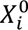, *X*_*i*_ refers to the coordinates of the ith atom in the reference structure and generated structure, respectively. *N* refers to the total atom number in the backbone of the protein. When calculating the RMSD, the protein alignment should be conducted to obtain the best-fit between the two structures in the Cartesian space. Here we use the MD analysis python package to calculate all RMSD values[36].

### 3.6 Jensen-Shannon (JS) divergence

To further evaluate the similarity between the generated protein structures and the reference structures from MD simulations, the Jensen-Shannon Divergence (JS Divergence) was used to compare the probability distributions of certain structural features, such as dihedral angles or bond angles[37,38]. JS divergence is a symmetric, information-theoretic measure that quantifies the similarity between two probability distributions.

Given two probability distributions P and Q over the same probability space, the JS Divergence is defined as a smoothed and symmetric version of KL Divergence. It measures the divergence between each distribution and its average. The JS Divergence is given by:

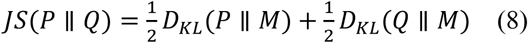

where 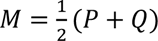 is the average of the two distributions, and *D_KL_* (*P* ∥ *M*) and *D*_KL_ (*Q* ∥ *M*) are the KL Divergences between the distributions and the mean distribution. The KL Divergence for two distributions P and Q is defined as:

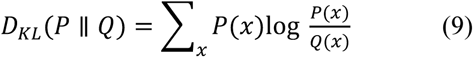

## 4. Discussion

In this study, we have developed a novel approach to protein structure generation by employing angular deviations as the data flow for a transformer-based diffusion model. This method has shown significant improvements in generating stable and accurate protein backbone structures compared to previous approaches.

One of the key innovations in our work is the use of angular deviations, rather than absolute backbone angles, as the data flow for our model. The motivation for this choice stems from the inherent SE(3) symmetry of protein structures, which describes the group of spatial transformations (rotations and translations) that do not alter the physical properties of the system. Previous work by Kevin E. Wu et al. demonstrated the utility of using six types of backbone angles (’ϕ’, ‘ψ’, ‘ω’, ‘θ₁’, ‘θ₂’, ‘θ₃’) as model inputs[27]. This approach effectively reduces the degrees of freedom from 9N (where N is the number of residues) to 6N, thereby lowering the dimensionality of the data and making the problem more tractable.

Moreover, we observed that the denoising process—where the model transitions from a disordered to an ordered state—was significantly more robust when using angular deviations. The generated structures moved from a disordered state_(figure 8a, timestep 0) to a partially folded intermediate (figure 9a, timestep 10), and then to a folded state_(figure 9a, timestep 50) more rapidly and reliably. This contrasts with the model trained on absolute angles, where the structures often pass through an unfolded state_(figure 9b, timestep 10), and then slowly to a partially folded intermediate_(figure9b, timestep 850), and then to a folded state_(figure9b, timestep 999), which increases the risk of misfolding at each intermediate step. Misfolding at any stage can prevent the final structure from achieving the correct fold, leading to a loss in structural fidelity. This difference in folding pathways underscores the importance of input representation in the accuracy and reliability of protein structure prediction (Figure 9). For more details see supplement material.

**Figure 9.**
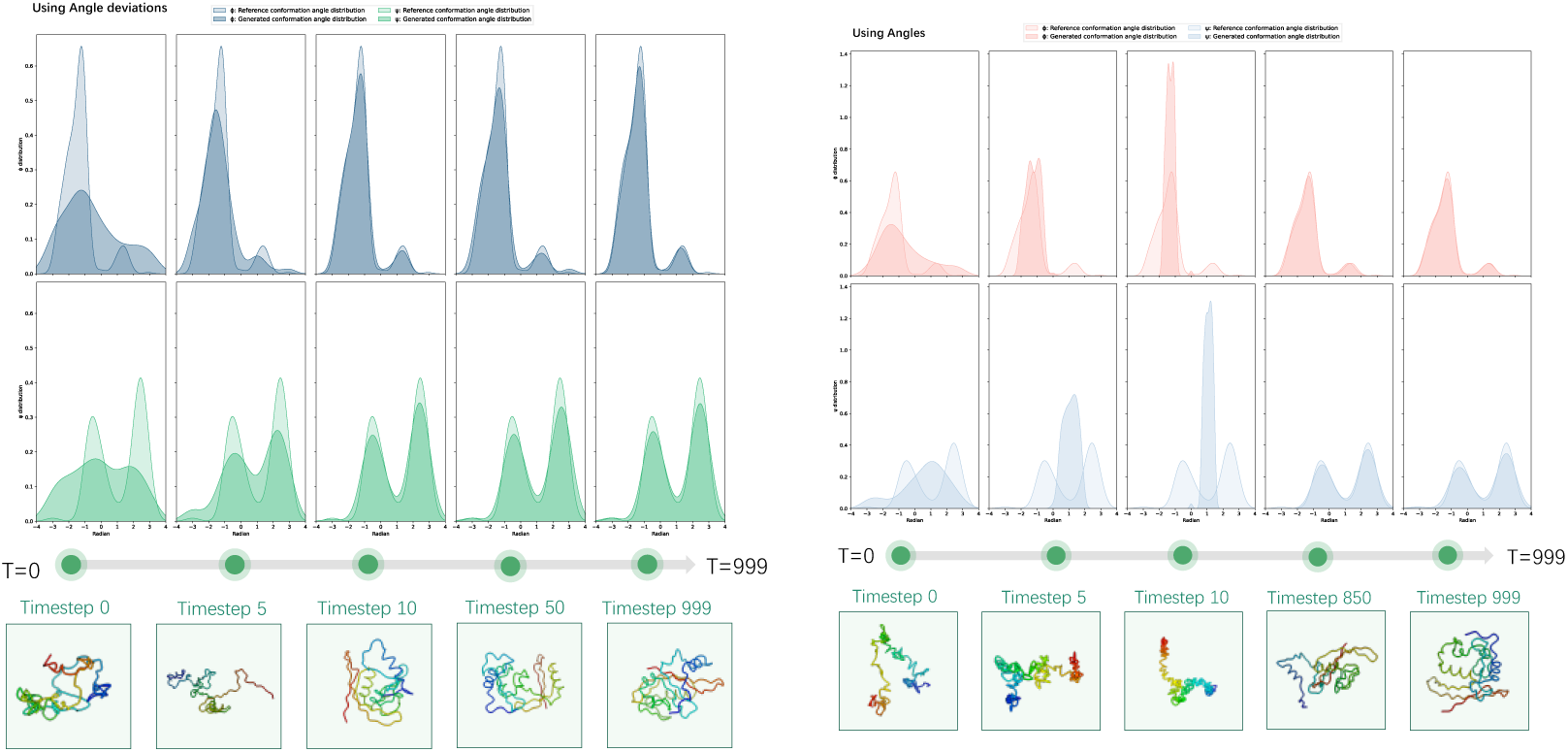
Denoising process of using angles and angle deviations for the Dark state system. Shifts in angle distributions, and conformations. (a) using angle deviations(b) using angles.

Furthermore, in protein simulation systems involving small molecules, the final equilibrium conformational ensemble is influenced by the interaction between the small molecules and the protein[39]. This suggests a limitation in current methods that generate protein conformational ensembles based solely on amino acid sequences, as they may not fully capture the impact of small molecule interactions on the final protein structure[21].

While our model has demonstrated the ability to generate high-quality protein structures, we observed that some of the generated structures exhibited atomic clashes, where interatomic distances were shorter than physically plausible limits[40]. These clashes result in non-physical configurations that would be unstable in a real biological system, to address this issue, we employed force field-based optimization, using PyRosetta[29], to refine the generated structures.

In future work, we aim to integrate force fields directly into our model to improve the quality of the generated structures. Additionally, The rapid advancements in video generation models present an exciting opportunity for the future of protein structure prediction[41–43]. Currently, our model generates static structures, capturing snapshots of protein conformations. However, proteins are dynamic entities, and their function is often tied to their conformational flexibility and the pathways they follow during folding. we envision training a model capable of generating entire trajectories, including both temporal and pathway information[44].

### Conclusion

In this study, we developed a transformer-based diffusion model that employs angular deviations to generate protein backbone structures with greater accuracy and efficiency. By utilizing the inherent SE(3) symmetry of protein structures and leveraging angular deviations from reference conformations, our model significantly outperforms traditional approaches that rely on absolute angle inputs. The validation experiments demonstrated the model’s capability to generate more ensembles with a large range of RMSD values comparable to those from MD simulations, while also revealing new conformations that extend beyond the training data. Moreover, the use of angular deviations resulted in a more stable and reliable folding process, as indicated by lower rates of misfolding and closer alignment with the dynamic behavior observed in MD simulations.

## Reference

1. Taylor, W.R. Exploring Protein Fold Space. Biomolecules 2020, 10, 193, doi:10.3390/biom10020193.

2. Pantsar, T. The Current Understanding of KRAS Protein Structure and Dynamics. Computational and Structural Biotechnology Journal 2020, 18, 189–198, doi:10.1016/j.csbj.2019.12.004.

3. Shukla, R.; Tripathi, T. Molecular Dynamics Simulation in Drug Discovery: Opportunities and Challenges. In Innovations and Implementations of Computer Aided Drug Discovery Strategies in Rational Drug Design; Singh, S.K., Ed.; Springer: Singapore, 2021; pp. 295–316 ISBN 9789811589362.

4. Shaw, D.E.; Adams, P.J.; Azaria, A.; Bank, J.A.; Batson, B.; Bell, A.; Bergdorf, M.; Bhatt, J.; Butts, J.A.; Correia, T.;, et al. Anton 3: Twenty Microseconds of Molecular Dynamics Simulation before Lunch. In Proceedings of the Proceedings of the International Conference for High Performance Computing, Networking, Storage and Analysis; Association for Computing Machinery: New York, NY, USA, November 13 2021; pp. 1–11.

5. Rosta, E.; Hummer, G. Error and Efficiency of Replica Exchange Molecular Dynamics Simulations. The Journal of Chemical Physics 2009, 131, 165102, doi:10.1063/1.3249608.

6. Wang, J.; Arantes, P.R.; Bhattarai, A.; Hsu, R.V.; Pawnikar, S.; Huang, Y.M.; Palermo, G.; Miao, Y. Gaussian Accelerated Molecular Dynamics: Principles and Applications. WIREs Comput Mol Sci 2021, 11, e1521, doi:10.1002/wcms.1521.

7. Zuo, K.; Kranjc, A.; Capelli, R.; Rossetti, G.; Nechushtai, R.; Carloni, P. Metadynamics Simulations of Ligands Binding to Protein Surfaces: A Novel Tool for Rational Drug Design. Phys. Chem. Chem. Phys. 2023, 25, 13819–13824, doi:10.1039/D3CP01388J.

8. Wang, J.; Peng, C.; Yu, Y.; Chen, Z.; Xu, Z.; Cai, T.; Shao, Q.; Shi, J.; Zhu, W. Exploring Conformational Change of Adenylate Kinase by Replica Exchange Molecular Dynamic Simulation. Biophysical Journal 2020, 118, 1009–1018, doi:10.1016/j.bpj.2020.01.001.

9. Tian, H.; Jiang, X.; Xiao, S.; La Force, H.; Larson, E.C.; Tao, P. LAST: Latent Space-Assisted Adaptive Sampling for Protein Trajectories. J. Chem. Inf. Model. 2023, 63, 67–75, doi:10.1021/acs.jcim.2c01213.

10. Joshi, S.Y.; Deshmukh, S.A. A Review of Advancements in Coarse-Grained Molecular Dynamics Simulations. Molecular Simulation 2021, 47, 786–803, doi:10.1080/08927022.2020.1828583.

11. Bandi, A.; Adapa, P.V.S.R.; Kuchi, Y.E.V.P.K. The Power of Generative AI: A Review of Requirements, Models, Input–Output Formats, Evaluation Metrics, and Challenges. Future Internet 2023, 15, 260, doi:10.3390/fi15080260.

12. Xiao, S.; Song, Z.; Tian, H.; Tao, P. Assessments of Variational Autoencoder in Protein Conformation Exploration. J Comput Biophys Chem 2023, 22, 489–501, doi:10.1142/s2737416523500217.

13. Zhu, J.-J.; Zhang, N.-J.; Wei, T.; Chen, H.-F. Enhancing Conformational Sampling for Intrinsically Disordered and Ordered Proteins by Variational Autoencoder. IJMS 2023, 24, 6896, doi:10.3390/ijms24086896.

14. Eguchi, R.R.; Choe, C.A.; Huang, P.-S. Ig-VAE: Generative Modeling of Protein Structure by Direct 3D Coordinate Generation. PLoS Comput Biol 2022, 18, e1010271, doi:10.1371/journal.pcbi.1010271.

15. Tian, H.; Jiang, X.; Trozzi, F.; Xiao, S.; Larson, E.C.; Tao, P. Explore Protein Conformational Space With Variational Autoencoder. Front. Mol. Biosci. 2021, 8, 781635, doi:10.3389/fmolb.2021.781635.

16. Zhu, J.; Li, Z.; Tong, H.; Lu, Z.; Zhang, N.; Wei, T.; Chen, H.-F. Phanto-IDP: Compact Model for Precise Intrinsically Disordered Protein Backbone Generation and Enhanced Sampling. Briefings in Bioinformatics 2023, 25, bbad429, doi:10.1093/bib/bbad429.

17. Anand, N.; Eguchi, R.; Huang, P.-S. Fully Differentiable Full-Atom Protein Backbone Generation. 2019.

18. Wang, Y.; Wang, L.; Shen, Y.; Wang, Y.; Yuan, H.; Wu, Y.; Gu, Q. Protein Conformation Generation via Force-Guided SE(3) Diffusion Models 2024.

19. Zheng, S.; He, J.; Liu, C.; Shi, Y.; Lu, Z.; Feng, W.; Ju, F.; Wang, J.; Zhu, J.; Min, Y.;, et al. Predicting Equilibrium Distributions for Molecular Systems with Deep Learning. Nat Mach Intell 2024, 6, 558–567, doi:10.1038/s42256-024-00837-3.

20. Watson, J.L.; Juergens, D.; Bennett, N.R.; Trippe, B.L.; Yim, J.; Eisenach, H.E.; Ahern, W.; Borst, A.J.; Ragotte, R.J.; Milles, L.F.;, et al. De Novo Design of Protein Structure and Function with RFdiffusion. Nature 2023, 620, 1089–1100, doi:10.1038/s41586-023-06415-8.

21. Jing, B.; Berger, B.; Jaakkola, T. AlphaFold Meets Flow Matching for Generating Protein Ensembles.; October 28 2023.

22. Tang, X.; Dai, H.; Knight, E.; Wu, F.; Li, Y.; Li, T.; Gerstein, M. A Survey of Generative AI for de Novo Drug Design: New Frontiers in Molecule and Protein Generation. Briefings in Bioinformatics 2024, 25, bbae338, doi:10.1093/bib/bbae338.

23. Vaidya, A.T.; Chen, C.-H.; Dunlap, J.C.; Loros, J.J.; Crane, B.R. Structure of a Light-Activated LOV Protein Dimer That Regulates Transcription. Sci Signal 2011, 4, ra50, doi:10.1126/scisignal.2001945.

24. Zoltowski, B.D.; Schwerdtfeger, C.; Widom, J.; Loros, J.J.; Bilwes, A.M.; Dunlap, J.C.; Crane, B.R. Conformational Switching in the Fungal Light Sensor Vivid. Science 2007, 316, 1054– 1057, doi:10.1126/science.1137128.

25. Zhou, H.; Zoltowski, B.D.; Tao, P. Revealing Hidden Conformational Space of LOV Protein VIVID Through Rigid Residue Scan Simulations. Sci Rep 2017, 7, 46626, doi:10.1038/srep46626.

26. Zhou, H.; Dong, Z.; Verkhivker, G.; Zoltowski, B.D.; Tao, P. Allosteric Mechanism of the Circadian Protein Vivid Resolved through Markov State Model and Machine Learning Analysis. PLoS Comput Biol 2019, 15, e1006801, doi:10.1371/journal.pcbi.1006801.

27. Wu, K.E.; Yang, K.K.; Van Den Berg, R.; Alamdari, S.; Zou, J.Y.; Lu, A.X.; Amini, A.P. Protein Structure Generation via Folding Diffusion. Nat Commun 2024, 15, 1059, doi:10.1038/s41467-024-45051-2.

28. Ho, J.; Jain, A.; Abbeel, P. Denoising Diffusion Probabilistic Models 2020.

29. Chaudhury, S.; Lyskov, S.; Gray, J.J. PyRosetta: A Script-Based Interface for Implementing Molecular Modeling Algorithms Using Rosetta. Bioinformatics 2010, 26, 689–691, doi:10.1093/bioinformatics/btq007.

30. Herrou, J.; Crosson, S. Function, Structure and Mechanism of Bacterial Photosensory LOV Proteins. Nat Rev Microbiol 2011, 9, 713–723, doi:10.1038/nrmicro2622.

31. Freddolino, P.L.; Gardner, K.H.; Schulten, K. Signaling Mechanisms of LOV Domains: New Insights from Molecular Dynamics Studies. Photochem Photobiol Sci 2013, 12, 1158–1170, doi:10.1039/c3pp25400c.

32. Jorgensen, W.L.; Chandrasekhar, J.; Madura, J.D.; Impey, R.W.; Klein, M.L. Comparison of Simple Potential Functions for Simulating Liquid Water. The Journal of Chemical Physics 1983, 79, 926–935, doi:10.1063/1.445869.

33. Devlin, J.; Chang, M.-W.; Lee, K.; Toutanova, K. BERT: Pre-Training of Deep Bidirectional Transformers for Language Understanding. In Proceedings of the Proceedings of the 2019 Conference of the North American Chapter of the Association for Computational Linguistics: Human Language Technologies, Volume 1 (Long and Short Papers); Burstein, J., Doran, C., Solorio, T., Eds.; Association for Computational Linguistics: Minneapolis, Minnesota, June 2019; pp. 4171–4186.

34. Kalra, D.S.; Barkeshli, M. Why Warmup the Learning Rate? Underlying Mechanisms and Improvements 2024.

35. Girshick, R. Fast R-CNN. In Proceedings of the 2015 IEEE International Conference on Computer Vision (ICCV); December 2015; pp. 1440–1448.

36. Gowers, R.; Linke, M.; Barnoud, J.; Reddy, T.; Melo, M.; Seyler, S.; Domański, J.; Dotson, D.; Buchoux, S.; Kenney, I.;, et al. MDAnalysis: A Python Package for the Rapid Analysis of Molecular Dynamics Simulations.; Austin, Texas, 2016; pp. 98–105.

37. McClendon, C.L.; Hua, L.; Barreiro, G.; Jacobson, M.P. Comparing Conformational Ensembles Using the Kullback–Leibler Divergence Expansion. J. Chem. Theory Comput. 2012, 8, 2115– 2126, doi:10.1021/ct300008d.

38. Fuglede, B.; Topsoe, F. Jensen-Shannon Divergence and Hilbert Space Embedding. In Proceedings of the International Symposium onInformation Theory, 2004. ISIT 2004. Proceedings.; June 2004; pp. 31-.

39. Protein–Small Molecule Interactions in Native Mass Spectrometry | Chemical Reviews Available online: https://pubs.acs.org/doi/full/10.1021/acs.chemrev.1c00293?casa_token=FdlISRwaNKgAAAAA%3AOY7S6WUG3RdWL_vcnJajtV7UDioMZ5MEEr6esygAui34uziMzkVCs9675-G_RMvYJkdbKogw3Kl3hXXq (accessed on 10 September 2024).

40. Lu, J.; Zhong, B.; Zhang, Z.; Tang, J. STR2STR: A SCORE-BASED FRAMEWORK FOR ZERO-SHOT PROTEIN CONFORMATION SAMPLING. 2024.

41. Yang, R.; Srivastava, P.; Mandt, S. Diffusion Probabilistic Modeling for Video Generation. Entropy 2023, 25, 1469, doi:10.3390/e25101469.

42. Liu, Y.; Cun, X.; Liu, X.; Wang, X.; Zhang, Y.; Chen, H.; Liu, Y.; Zeng, T.; Chan, R.; Shan, Y. EvalCrafter: Benchmarking and Evaluating Large Video Generation Models.; 2024; pp. 22139–22149.

43. Ruan, L.; Ma, Y.; Yang, H.; He, H.; Liu, B.; Fu, J.; Yuan, N.J.; Jin, Q.; Guo, B. MM-Diffusion: Learning Multi-Modal Diffusion Models for Joint Audio and Video Generation. In Proceedings of the 2023 IEEE/CVF Conference on Computer Vision and Pattern Recognition (CVPR); IEEE: Vancouver, BC, Canada, June 2023; pp. 10219–10228.

44. Kim, M.K.; Jernigan, R.L.; Chirikjian, G.S. Efficient Generation of Feasible Pathways for Protein Conformational Transitions. Biophysical Journal 2002, 83, 1620–1630, doi:10.1016/S0006-3495(02)73931-3.

