## Supplement material for "Angular Deviation Diffuser: A Transformer-Based Diffusion Model for Efficient Protein Conformational Ensemble Generation"

### Supplement materials

1 The Vivid (VVD) photoreceptor and LOV domain

The Vivid (VVD) photoreceptor is a key protein involved in regulating the circadian rhythms of the filamentous fungus *Neurospora crassa*. VVD is a member of the light, oxygen, or voltage (LOV) domain family, a group of small, modular protein domains widely conserved across various kingdoms of life, including bacteria, archaea, fungi, and plants. LOV domains mediate responses to blue light by inducing conformational changes in their associated proteins. This ability to sense and respond to environmental light conditions is fundamental for processes such as phototropism, circadian rhythm regulation, and photomorphogenesis[1,2].

LOV domains are structurally characterized by their binding to flavin-based cofactors—specifically flavin adenine dinucleotide (FAD), flavin mononucleotide (FMN), or riboflavin. These cofactors are essential for the blue light-induced photochemistry that drives the structural changes in LOV proteins. Upon blue-light absorption, the flavin cofactor undergoes a photoreduction, leading to the formation of a covalent bond with a conserved cysteine residue within the LOV domain. This triggers conformational shifts that propagate through the protein, resulting in downstream signaling events[3].

2 Molecular Dynamics Simulation

In this study, we employed molecular dynamics (MD) simulations to investigate the conformational changes in VVD between its dark-adapted and light-activated states. The initial structures for the MD simulations were obtained from the crystal structures of VVD in its dark state (PDB ID: 2PD7) and light state (PDB ID: 3RH8), both of which include the FAD cofactor.

To ensure consistency in the sequence lengths between the dark and light states, we removed the 36th residue from the dark state structure. Additionally, given the biological similarity between FMN and FAD, we modified the FAD molecule in the crystal structures by removing the AMP moiety to generate an FMN-like molecule. The force field parameters for FMN were adopted from previous work[3], allowing us to model its behavior accurately within the simulation system.

A total of two simulation systems were constructed based on the modified crystal structures. Hydrogen atoms were added to each structure, and the systems were solvated using the explicit water model TIP3P[4]. The systems were then neutralized by adding sodium cations and chloride anions.

Initially, we performed 10 nanoseconds (ns) of isothermal-isobaric ensemble (NPT) MD simulations to equilibrate the systems. Following this, 1.1 microseconds (μs) of canonical ensemble (NVT) Langevin MD simulations were conducted at a temperature of 300 K. The first 100 ns of these simulations were discarded to ensure proper equilibration, and the subsequent 1 μs of simulation data was utilized for analysis. The total number of TIP3P water molecules added to the simulation systems was 7,240 and 9429 for the dark and light configurations, respectively.

For all simulations, the SHAKE algorithm[5] was employed to constrain bonds involving hydrogen atoms, allowing for a time step of 2 femtoseconds (fs). Simulation trajectories were saved at intervals of 100 picoseconds (ps). A cubic simulation box and periodic boundary conditions were applied in all cases. Electrostatic interactions were calculated using the particle mesh Ewald (PME) method, ensuring accurate long-range interaction calculations[6].

All MD simulations were carried out using the CHARMM simulation package version 41b1[7], with support for graphics processing unit (GPU) acceleration based on OpenMM, significantly enhancing computational efficiency[8].

From these simulations, 6001 protein conformations have been saved to be the dataset in each system(Figure S1)

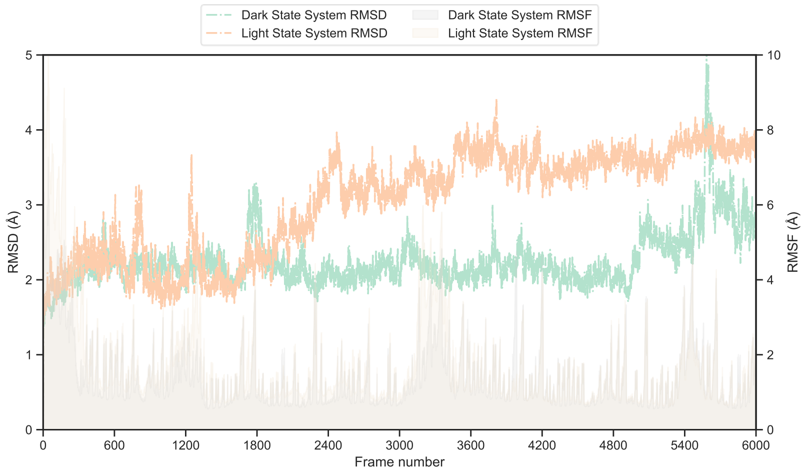

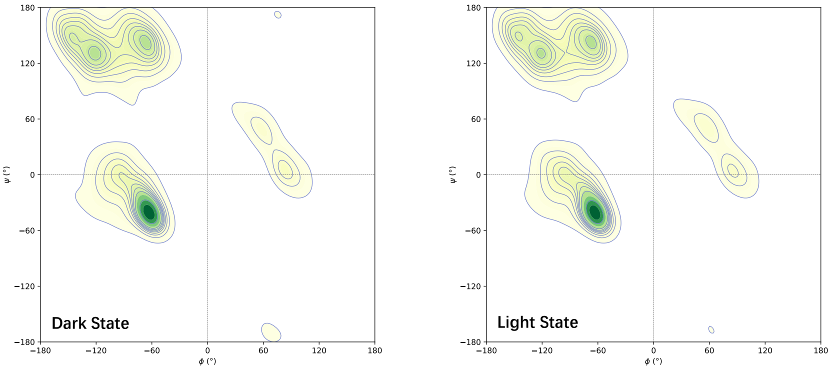


**Figure S1 Dark and Light state MD simulations.** (a) RMSD regarding the first frame and RMSF regarding the averaged conformation for each system (b) Ramachandra plot for each system

3 Transformer model

The transformer model in this work is based on a modified BERT architecture, designed to handle angle deviation data[9]. It uses a hidden size of 384 with 12 transformer layers and 12 attention heads. The intermediate size in the feed-forward network is set to 768, and a dropout rate of 0.1 is applied to prevent overfitting. The model incorporates relative positional embeddings to capture sequence order and supports input sequences up to a maximum length of 147. Ange deviation with diffusion time information is embedded using random Fourier features, ensuring that the model can capture time-based dependencies effectively.

**TableS1** The hyperparameters in our model

| **Hyperparameter** | **Value** |
| --- | --- |
| Hidden Size | 384 |
| Number of Layers | 12 |
| Number of Attention Heads | 12 |
| Intermediate Size | 768 |
| Dropout Rate | 0.1 |
| Max Position Embeddings | 147 |
| Position Embedding Type | Relative Key |
| Temporal Embedding Type | Random Fourier Features |

4 Refinement

In this study, we utilized *PyRosetta* to reconstruct and refine side-chain conformations for backbone conformations generated by our model[10]. *PyRosetta* integrates energy minimization with constraint-based optimization strategies, where the scoring function is critical in evaluating the energy landscape of a given protein conformation. This scoring function incorporates various energetic components, including van der Waals interactions, electrostatic forces, and hydrogen bonding.

To preserve structural integrity during the refinement process, coordinate constraints are applied, often employing harmonic potentials to restrict deviations from the initial atomic positions. The FastRelax protocol is a key refinement tool to achieve global protein structure optimization. By gradually reducing the strength of the constraints while minimizing the energy, the protocol enables the exploration of lower-energy conformations while maintaining proximity to the original structure.

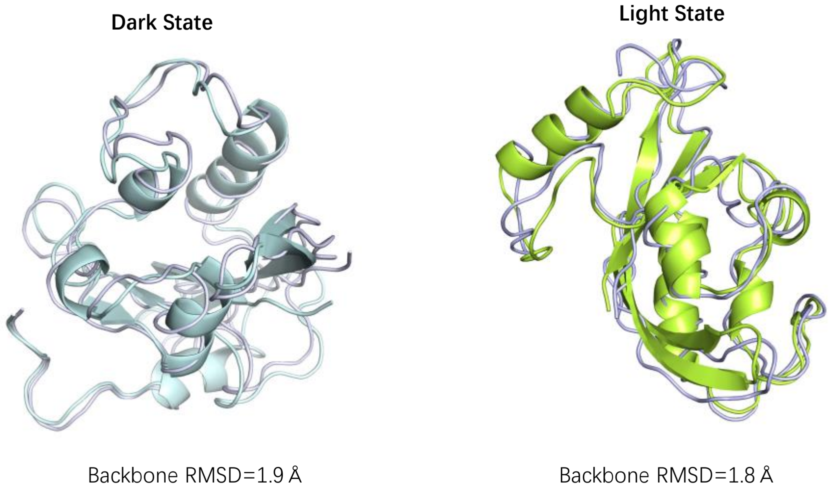


**Figure S2 Comparison of the directly generated conformations to refined ones through RMSD. Light-colored structure corresponds to conformation before refinement, while dark-colored structure represents conformation with refinement.**

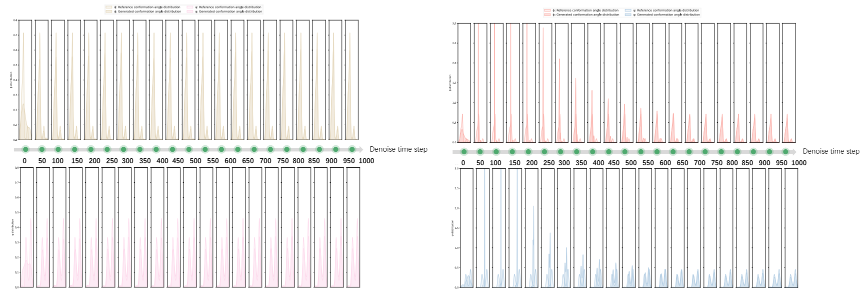


**Figure S3 The angle distribution changes in the denoise process for the Dark state.** (a) using absolute angles as data flow (b) using angle deviation as data flow

Reference


1. Zoltowski, B.D.; Schwerdtfeger, C.; Widom, J.; Loros, J.J.; Bilwes, A.M.; Dunlap, J.C.; Crane, B.R. Conformational Switching in the Fungal Light Sensor Vivid. *Science* **2007**, *316*, 1054–1057, doi:10.1126/science.1137128.
2. Vaidya, A.T.; Chen, C.-H.; Dunlap, J.C.; Loros, J.J.; Crane, B.R. Structure of a Light-Activated LOV Protein Dimer That Regulates Transcription. *Sci Signal* **2011**, *4*, ra50, doi:10.1126/scisignal.2001945.
3. Freddolino, P.L.; Gardner, K.H.; Schulten, K. Signaling Mechanisms of LOV Domains: New Insights from Molecular Dynamics Studies. *Photochem Photobiol Sci* **2013**, *12*, 1158–1170, doi:10.1039/c3pp25400c.
4. Jorgensen, W.L.; Chandrasekhar, J.; Madura, J.D.; Impey, R.W.; Klein, M.L. Comparison of Simple Potential Functions for Simulating Liquid Water. *The Journal of Chemical Physics* **1983**, *79*, 926–935, doi:10.1063/1.445869.
5. Kräutler, V.; van Gunsteren, W.F.; Hünenberger, P.H. A Fast SHAKE Algorithm to Solve Distance Constraint Equations for Small Molecules in Molecular Dynamics Simulations. *Journal of Computational Chemistry* **2001**, *22*, 501–508, doi:10.1002/1096-987X(20010415)22:5<501::AID-JCC1021>3.0.CO;2-V.
6. Essmann, U.; Perera, L.; Berkowitz, M.; Darden, T.; Lee, H.; Pedersen, L. A Smooth Particle Mesh Ewald Method. *J. Chem. Phys.* **1995**, *103*, 8577, doi:10.1063/1.470117.
7. Brooks, B.R.; Brooks, C.L.; MacKerell, A.D.; Nilsson, L.; Petrella, R.J.; Roux, B.; Won, Y.; Archontis, G.; Bartels, C.; Boresch, S.; et al. CHARMM: The Biomolecular Simulation Program. *J Comput Chem* **2009**, *30*, 1545–1614, doi:10.1002/jcc.21287.
8. Eastman, P.; Pande, V.S. OpenMM: A Hardware Independent Framework for Molecular Simulations. *Comput Sci Eng* **2015**, *12*, 34–39, doi:10.1109/MCSE.2010.27.
9. Devlin, J.; Chang, M.-W.; Lee, K.; Toutanova, K. BERT: Pre-Training of Deep Bidirectional Transformers for Language Understanding. In Proceedings of the Proceedings of the 2019 Conference of the North American Chapter of the Association for Computational Linguistics: Human Language Technologies, Volume 1 (Long and Short Papers); Burstein, J., Doran, C., Solorio, T., Eds.; Association for Computational Linguistics: Minneapolis, Minnesota, June 2019; pp. 4171–4186.

10. Chaudhury, S.; Lyskov, S.; Gray, J.J. PyRosetta: A Script-Based Interface for Implementing Molecular Modeling Algorithms Using Rosetta. *Bioinformatics* **2010**, *26*, 689–691, doi:10.1093/bioinformatics/btq007.
